# Is testing the bee-all and end-all of *varroa* eradication?

**DOI:** 10.1101/2024.07.18.604034

**Authors:** Isobel R. Abell, Jennifer A. Flegg, Thao P. Le, Christopher M. Baker

## Abstract

*Varroa destructor* is a significant European honeybee pest, impacting agricultural industries globally. Since arriving in 2022, Australia faces the possibility that *varroa* will become established in European honeybee colonies nationally. Australia initially pursued a strategy of testing and subsequently eliminating hives infested with *varroa*. These management efforts raise interesting questions about the interplay between hive testing and elimination, and the spread of *varroa* between hives. This study uses mathematical modelling to investigate how combined hive testing and elimination strategies impact the spread of *varroa* through a network of European honeybee hives. We develop a model of both within-hive reproduction of *varroa* and hive testing, and between-hive movement of *varroa* on a network of hives. This model is used to assess the impact of various testing and hive elimination strategies on the total number of hives eliminated on the network of hives. Each model simulation starts with a single infested hive, and from this we observed one of two dynamics: either the infestation is caught before spreading, or *varroa* spreads extensively through the network before being caught by testing. Within our model we implement two common hive testing methods – sugar shake and alcohol testing. A shared limitation of these testing methods is that they can only detect mites in a specific stage of their lifecycle. As such, testing is not only dependent on how many *varroa* mites are in a hive, but what lifecycle stage the mites are in at the time of testing. By varying testing and movement parameters, we see that this testing limitation greatly impacts the number of hives eliminated in various scenarios. We find that there are largely two invasion possibilities: either there is only a small incursion, or that *varroa* achieves complete spread on the network. Furthermore, testing earlier, or testing more frequently, does not guarantee a smaller invasion. Our model results suggest irregular testing schedules, e.g. testing multiple times in close succession rather than just once in a given timeframe, may help overcome the limitations of common hive testing strategies.

## 1 Introduction

Invasive pests and diseases cause large economic losses globally (Diagne et al. 2020). Designing and implementing effective control strategies to eliminate or, where this is not possible, manage invasive pests is crucial to maintain functioning agricultural sectors. *Varroa destructor* (henceforth referred to as *varroa*) is one of the most destructive parasites for European honeybees. The combination of physiological damage due to parasitism and the introduction of diseases carried by the mite, such as Deformed Wing Virus, has resulted in major colony losses for European honeybees globally (De La Rúa et al. 2009; Roberts, Anderson, and Durr 2017). Maintaining control over *varroa* infestations in European honeybee colonies presents a key challenge for beekeeping in Australia and around the world.

Historically, Australia has remained free of *varroa*. However, the abundance of European honeybees in Australia has wide reaching implications for both European honeybee and native bee colonies should *varroa* become established. In Australia, *Apis Mellifera* (European honeybees) are an invasive species, first introduced by English colonists around the 1800s (Paini 2004). Since then, they have remained in Australia both in managed colonies and in feral colonies, with Australia having one of the highest densities of feral European honeybees globally (Cunningham et al. 2022). Due to a reliance on European agricultural practises, European honeybees (both managed and feral) play a crucial role in supporting Australian agriculture. Furthermore, while native Australian bees can not be parasitised by *varroa*, they can contract diseases from European honeybees which have been parasitised by *varroa*. The introduction of diseases such as Deformed Wing Virus carried by *varroa* to native bees could result in high colony losses (Brettell et al. 2020). Thus, the establishment of *varroa* in Australia European honeybee colonies could lead to losses in a wide variety of agricultural sectors while also impacting native bee populations.

Following the latest detection of *varroa* in 2022, Australia has been faced with the potential establishment of *varroa* among European honeybee colonies nationally. While there have been previous detections of *varroa jacobsoni* (closely related, but distinct from *varroa destructor*) in Townsville, Queensland in 2016 and 2019, these outbreaks were declared eradicated (Australian Government 2024). In June 2022, *varroa destructor* was detected by sentinel hives in Port of Newcastle, New South Wales. The New South Wales Department of Primary Industries (NSW DPI) subsequently implemented an eradication strategy. This strategy was comprised of movement restrictions and euthanasia of infested hives in designated zones (New South Wales Department of Primary Industries 2022). However, in September 2023, NSW DPI transitioned to a management strategy following the unsuccessful eradication of *varroa*. This management strategy involved distributing miticide strips to treat infestations in individual hives and maintained movement restrictions in designated zones (Primary Industries 2023; New South Wales Department of Primary Industries 2024). States and territories around Australia, particularly those neighbouring New South Wales, have implemented surveillance and biosecurity measures to prevent *varroa* crossing state borders (Agriculture Victoria 2024; Department of Primary Industries and Regions 2022; Queensland Government 2024). While historically Australia has focused on biosecurity measures to remain *varroa*-free, recent incursions have forced Australia to implement eradication and management strategies to protect European honeybee colonies nationwide.

Mathematical modelling can be a useful tool to understand the dynamics of phenomena such as the spread of a pest or disease and the impact of different management strategies. In particular, network modelling can provide a framework to quantify the spread of a parasite or disease of a network of premises such as hives. Network models have been used in fields such as ecology, disease management and bushfire management to understand the spread of a phenomena through a network (Arancibia and Morin 2022; Firestone et al. 2011; Jiang et al. 2022). These types of models have also been used to study the spread of *varroa* and diseases carried by *varroa* through hives of European honeybees (Messan et al. 2017; Eberl and Muhammad 2022; Betti and Shaw 2021; Ibrahim and Dénes 2022). However, these studies have often focused on exploring how *varroa* or diseases travel through hives in a single apiary rather than through a naive population. In an Australian context, Owen et al. study the potential spread of varroa mite following importation in Victoria, Australia (Owen, Stevenson, and Scheerlinck 2021). This studied how far *varroa* could spread in an naive population before interventions, such as testing, are applied. Given the recent detection of *varroa* and subsequent eradication attempts, mathematical modelling is a valuable tool to assess the key drivers of *varroa* spread and the impact of combined testing and hive elimination strategies.

In this study, we model the spread of *varroa* on a network of hives under an elimination outbreak management strategy. We assume a strategy where all hives on a network are tested regularly and where hives with a positive detection of *varroa* are destroyed. Our modelling comprises of a within-hive model, incorporating reproduction and testing of *varroa* in a single hive, and a between-hive model, incorporating movement of *varroa* between hives. We assume an initially *varroa*-naive population, seeding infestations in each simulation in a single random hive. We vary the movement of *varroa* between hives, time between testing, testing start day, and test type, and assess their impact on the number of hives infested. Our work shows how modelling can be used to investigate the dynamics of *varroa* spread and assess the impact of various testing strategies in combination with hive elimination.

## 2 Methods

We model the spread of *varroa* through a network of hives by incorporating a) the reproduction and movement of *varroa*, and b) the testing and elimination of hives as part of an outbreak response. The model consists of 2 components: a within-hive model (Section 2.1), capturing the dynamics of mite reproduction and testing for a single hive; and a between-hive model (Section 2.2), capturing the dynamics of *varroa* movement between hives. Figure 1 presents an overview of the within-hive and between-hive model structures.

**Figure 1:**
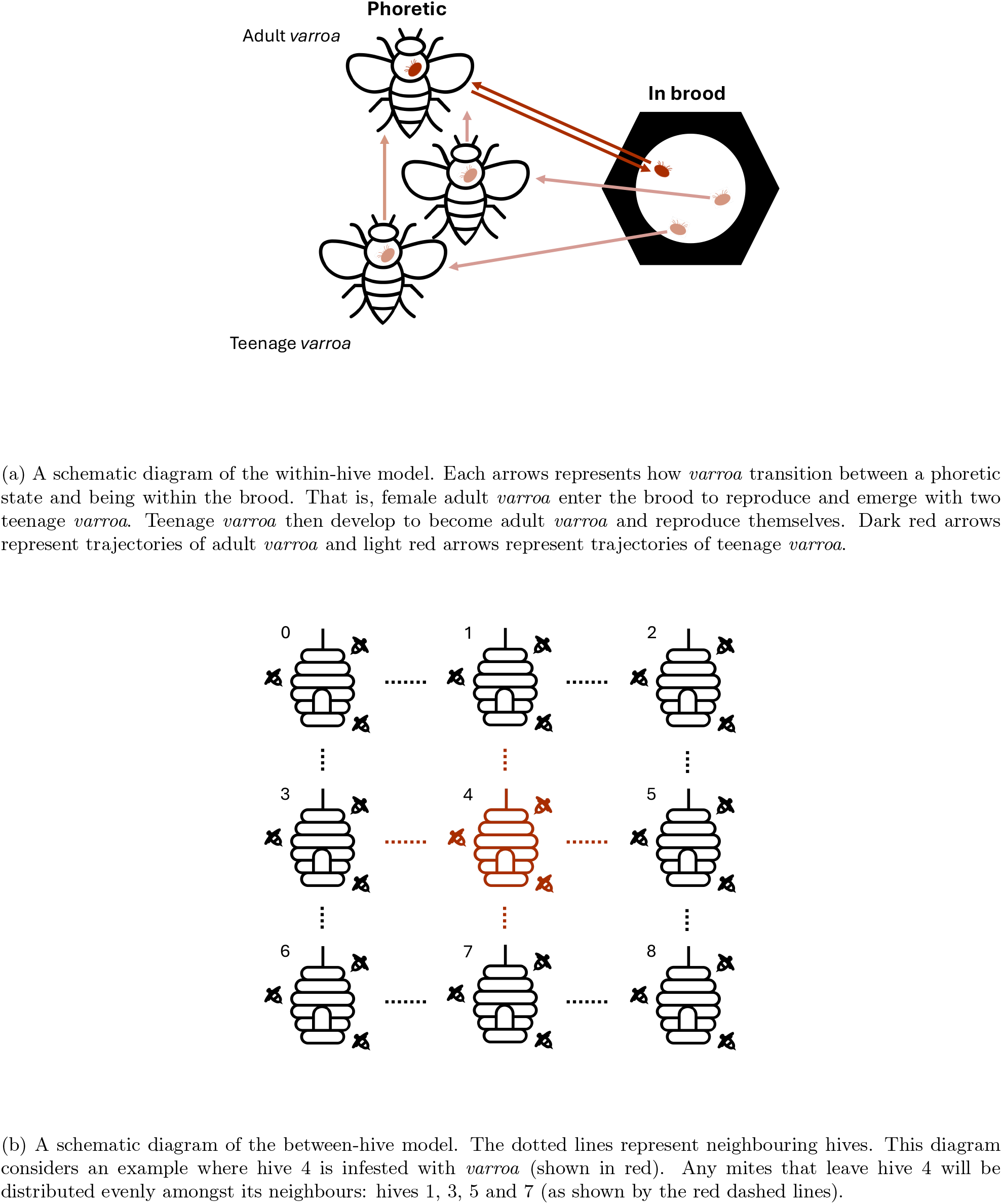
An overview of the model structure for both within-hive (a) and between-hive (b) models.

### 2.1 Within-hive model

The within-hive model tracks the total number of mites in each stage of a mite’s life-cycle. Testing, while occurring simultaneously across the network of hives at regular intervals, is implemented in each individual hive.

#### 2.1.1 Mite reproduction

The reproductive cycle of *varroa* is closely tied to that of European honeybees.

Before parasitising European honeybees, *varroa* co-evolved alongside the Asian honeybee (*Apis cerana*) (Rosenkranz, Aumeier, and Ziegelmann 2010). As such, the *varroa* reproductive cycle is closely tied to that of Asian and European honeybees. While there may be some variation in the *varroa* reproductive cycle in reality, we approximate it here with a deterministic model.

We incorporate the following dynamics from the *varroa* reproductive cycle into our model:

- *Varroa* spend their life in two states: a phoretic state and within sealed brood cells (cells in the hives within which bee larvae develop) as shown in Figure 1a. While phoretic, adult *varroa* can be transported both to brood cells (where they can reproduce) and between hives (to potentially seed a new infestation) (Rosenkranz, Aumeier, and Ziegelmann 2010). *Varroa* can spend between 5 and 11 days in a phoretic state (Huang 2019). Once *varroa* make it into a sealed brood cell they can reproduce (Rosenkranz, Aumeier, and Ziegelmann 2010).
- Upon entering the brood, a female *varroa* mite will lay both female and male eggs. These mites will mate with each other, resulting in the fertilisation of the daughter mites. Male mites die in the brood while mother and daughter mites emerge (fertilised) when the brood is uncapped (Rosenkranz, Aumeier, and Ziegelmann 2010). *Varroa* preferentially enter drone brood as they are capped for longer than worker brood (14 days) (Mortensen, Schmehl, and Ellis 2013) and can produce more daughter mites (2–3) (Rosenkranz, Aumeier, and Ziegelmann 2010).
- Daughter mites continue to develop outside the brood and will start to lay eggs 14 days after first emerging (BeeAware 2022).
- On average, adult mites will enter the brood to reproduce a total of 3 times before dying (Rosenkranz, Aumeier, and Ziegelmann 2010).

Notably, female mites emerge from the brood fertilised. This means a *varroa* infestation can start from a single imported mite.

Our model counts the number of female mites in each stage of the *varroa* lifecycle. Based on the above information, we assume a deterministic model of *varroa* reproduction with the following parameters:

- Adult mites go through 3 reproductive cycles before dying.
- Adult mites spend 5 days phoretic before entering the brood.
- Adult mites spend 14 days in the brood.
- Adult mites produce 2 teenage mites when reproducing in the brood.
- Teenage mites spend 14 days phoretic before developing into adult mites.

#### 2.1.2 Mite death

There are two ways individual mites can die in our model: through coming to the end of their lifecycle and through accidentally falling off a bee. The first type of death is deterministic, mites that have gone through three reproductive cycles will be assumed to die. The second type is stochastic, phoretic mites have a chance of falling off a bee to the bottom of the hive where they will die (Rosenkranz, Aumeier, and Ziegelmann 2010). In our model, we assume mites have a 5% chance of falling off bees over their lifetime. Full implementation details can be found in the supplementary material.

#### 2.1.3 Testing hives for *varroa* infestation

Our modelling considers a combined testing and hive elimination strategy to manage a *varroa* outbreak across a network of hives. In each simulation, each hive is tested for *varroa* infestation on given testing days. When these testing days occur is determined by how long after the first incursion testing starts and how much time passes between testing.

We implement two types of testing common among Australian beekeepers: alcohol wash and sugar shake testing methods (Agriculture Victoria 2023b). Both tests sample a small number of worker bees (∼ 300 bees) to see if they are carrying *varroa* and thus can only capture phoretic *varroa* (Frost 2023). We therefore compare the testing methods by considering the differing proportion of mites removed from the sampled bees by each method.

We define a positive test result as at least one *varroa* mite is found during testing. We assume testing as 100% specificity, i.e if a mite is found during testing it is *varroa* and not a different species of mite. We assume 300 bees are sampled during testing (Frost 2023). The probability of detecting one mite on one sampled bee is given by:

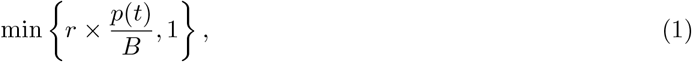

where *r* denotes the proportion of mites removed from sampled bees by the considered test method (akin to test sensitivity), *p*(*t*) denotes the total number of phoretic *varroa* in the considered hive at a given time *t* and *B* denotes the total number of bees in a hive (assumed to be constant over time). If *p*(*t*) < *B*, we assume there will be at most one phoretic *varroa* on each bee. If *p*(*t*) > *B*, we assume there are multiple phoretic *varroa* on each bee, i.e. every bee carries at least one mite. When calculating the probability of detecting one mite on one sampled bee, we take a minimum with 1 to ensure this probability is at most 1. We can use Eq. (1) to calculate the probability of detecting no *varroa* on one sampled bee:

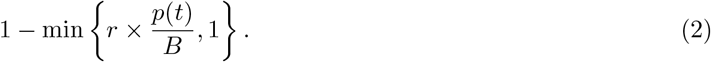

Given Eq. (2), we can calculate the probability of detecting no *varroa* mites on any sampled bees:

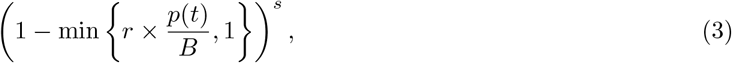

Where *s* = 300 denotes the total number of bees sampled in the testing process. We can use Eq. 3 to calculate the probability of detecting at least one *varroa* on the sampled bees:

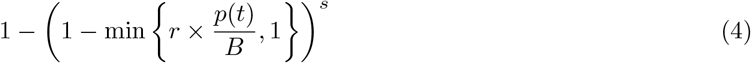

In our modelling, we assume the total number of bees in a hive is constant at 40,000 (New South Wales Department of Primary Industries 2020) and 300 bees are sampled during testing (*B* = 40000 and *s* = 300). We assume alcohol wash removes 70% of mites on sampled bees and sugar shake only 40% (Agriculture Victoria 2023a; Flores, Gil, and Padilla 2015). This gives removal proportions of *r* = 0.7 for alcohol wash and *r* = 0.4 for sugar shake. Therefore, the probability of returning a positive test result for both test methods is given by:

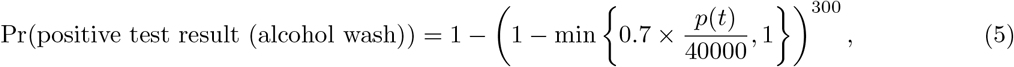

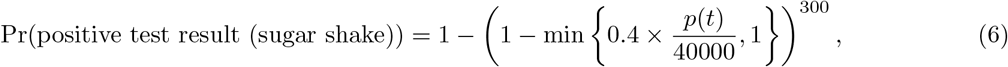

where *p*(*t*) is calculated from the reproduction mathematical model.

Eqs. (5) and (6) can be equivalently described using the binomial distribution:

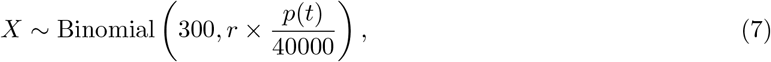

where Eq. (5) is equivalently the probability *X* ≥ 1 given *r* = 0.7, and Eq. (6) the probability *X* ≥ 1 given *r* = 0.4.

In our model, on a given testing day a uniform random number is drawn for each hive. If the random number is less than the probability of a positive test result, *varroa* has been detected in the hive and it will be eliminated on the same day.

#### 2.1.4 Hive elimination

In our assumed outbreak management strategy, if a hive returns a positive test result on a testing day, it is eliminated. That is, all bees are killed and all equipment sterilised, thus killing all mites in the hive. In our model, when a positive test result is returned, all mites in the hive are killed and the hive is designated as ‘dead’. That is, the number of *varroa* in each lifecycle stage is set to 0 for a dead hive. As equipment sterilisation usually results in the destruction of hive equipment, we assume if a hive has been eliminated it can no longer receive any mites from its neighbours.

### 2.2 Between-hive model

The between-hive model uses the number of phoretic *varroa* in each hive (as determined by the within-hive model) to calculate the number of mites moving between hives.

#### 2.2.1 Hive neighbourhoods

To define movement of mites between hives, we must first define the neighbourhoods of each hive. For each simulation, we assume there are 9 hives on a 3 *×* 3 grid (as demonstrated in Figure 1b). The total number of hives has been chosen arbitrarily for this study. Any grid size can be used, however computational time for our modelling greatly increases with grid size. Each hive’s neighbours are defined as the adjacent hives on the grid, i.e. the hive above, below, to the left and to the right (given there are hives in these positions). If a hive has previously been eliminated due to detection of *varroa*, it can still be a neighbour of an adjacent hive, but cannot receive any *varroa* through between-hive movement.

#### 2.2.2 Movement between hives

Bees can move between hives for a number of reasons, for example to steal resources from a weak neighbouring hive (called robbing behaviour), or because they drifted into a new hive on the way home (Messan et al. 2017). If a bee moving into a new hive is carrying any *varroa*, they can unwittingly seed an infestation in this new hive. In our model, the movement of bees (and the mites they may be carrying) is implemented stochastically. That is, every day we sample from a distribution to determine the number of *varroa* leaving a hive and entering neighbouring hives.

On average, about a third of a European honeybee colony leaves the hive to forage (Garvey 2013). In our model, we assume every hive has 40,000 bees and that a third of these bees leave the hive on a given day. Of the bees that leave the hive, we assume that a defined proportion (denoted *α* where 0 < *α* < 1) do not make it home to their original hive and migrate to a neighbouring hive. The number of bees migrating from one hive to a neighbouring hive each day is therefore given by:

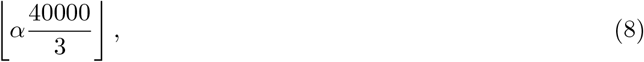

where we consider the floor to ensure an integer number of bees.

Given the number of bees migrating to a new hive, we can then calculate the number of *varroa* leaving a hive. We assume that if there are fewer phoretic mites than bees in a hive, if a bee carries a mite it will carry at most one. If there are more phoretic mites than bees in a hive, we assume every bee carries at least one mite. To calculate the number of mites leaving a hive, we sample from a hypergeometric distribution. We define a draw as a bee migrating between hives, and a success as the state where there is a *varroa* on that bee. As such, we assume a population size of 40,000 (total bees), the number of success states as the number of phoretic mites on a given day (*p*(*t*)), and the number of draws as the number of bees migrating from one hive to another (as defined by Eq. (8)).

For every day in our simulation, the model randomly samples from a hypergeometric distribution with the previously defined parameters to determine the number of mites leaving each hive in the network. Once the number of mites leaving a hive is calculated, they are equally distributed among the neighbouring hives. If a hive has been eliminated previously, it cannot receive any mites from its neighbours.

### 2.3 Simulating an infestation

For each model simulation, we seed an infestation in a single hive with three mites. Each simulations runs through time until there are no mites in any of the hives.

The initially infested hive is chosen at random. When we consider a grid of nine hives, the number of neighbours each hive has varies across the grid. That is: Hives 0, 2, 6 and 8 have two neighbours, Hives 1, 3, 5 and 7 have three neighbours and Hive 4 has four neighbours. When *varroa* move from one hive to its neighbours, we assume the mites are equally distributed among its neighbours. Where the outbreak is seeded and how many neighbours the initial hive has will therefore impact the total spread of *varroa*. To reduce the impact of these effects on our collated results, we randomly choose an initially infested hive in each simulation.

Each simulation is seeded by placing three *varroa* in random (phoretic) points of the adult reproductive cycle in the initially infested hive. As mite reproduction is deterministic, by varying the lifecycle stage of the initial mites we ensure that the mites are not entering and exiting the brood and reproducing in sync with one another across all hives.

Further modelling details, along with modelling code, can be found in the supplementary material.

## 3 Results

The results are structured into three sections: 3.1 Testing dynamics, 3.2 Emergent dynamics, and 3.3 Sensitivity analysis. Section 3.3 is further split into three subsections: 3.3.1 Movement between hives, 3.3.2 Test timing, and 3.3.3 Mites removed by testing.

All results consider the total number of hives eliminated by the end of the outbreak. That is, the number of hives that tested positive for *varroa* infestation and were subsequently eliminated. Each model simulation runs until there are no mites in any hive. This is due to a hive being (a) never infested, (b) infested, but the mites have died out before the hive tests positive for *varroa*, or (c) the hive tested positive for *varroa* and was eliminated. As we consider a total of nine hives, the total number of hives eliminated can only be between 0 hives and 9 hives.

Unless otherwise stated, we assume default modelling parameters as described in Table 1.

**Table 1:**
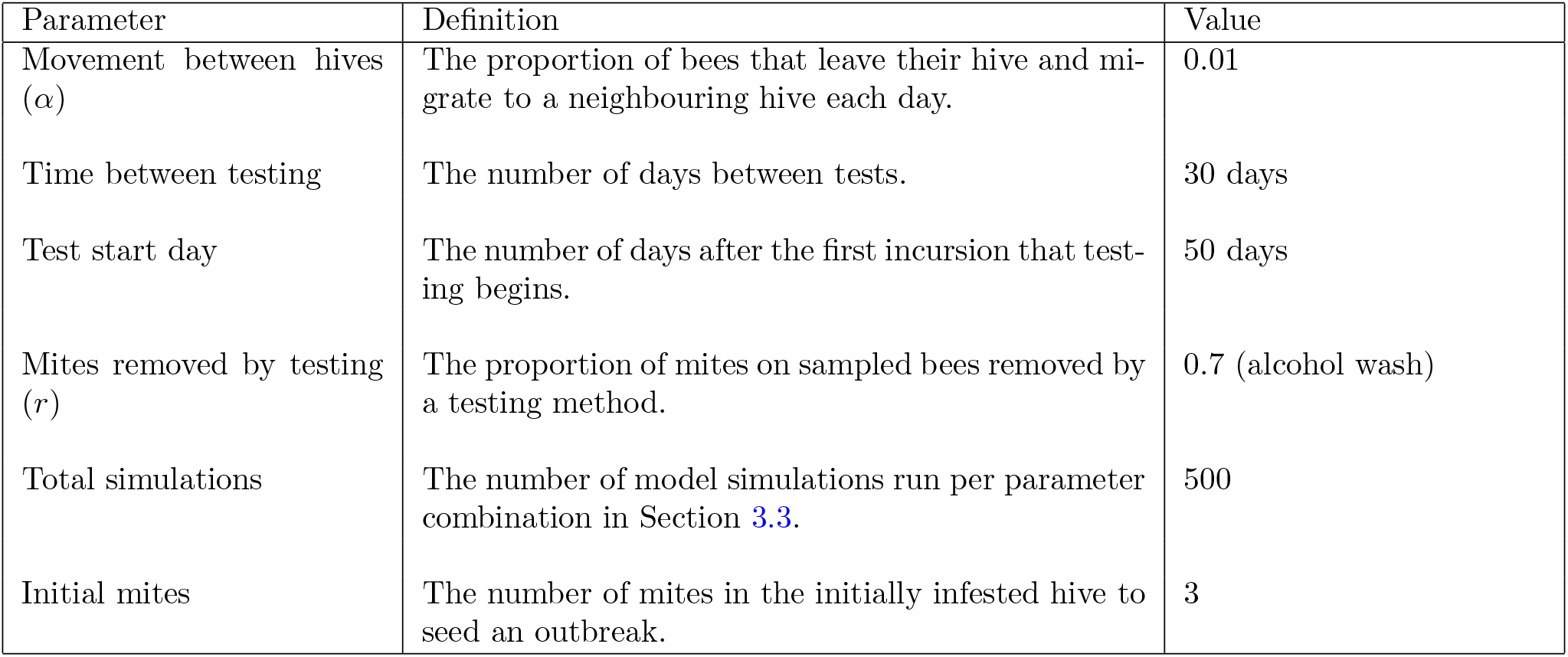
Default model parameter values.

### 3.1 Testing dynamics

When testing a hive for *varroa* infestation, the probability of positive detection depends on whether the testing strategy considers all mites in a hive or only phoretic mites. In this study, we only consider sugar shake and alcohol wash testing methods, the two most commonly used by beekeepers in Australia. Both methods aim to dislodge mites from sampled bees, and therefore can only ever capture phoretic *varroa*. The probability of positive detection for these tests is therefore dependent on the proportion of mites dislodged by testing, the number of bees sampled and the number of phoretic *varroa* in a hive.

Figure 2 shows an example of how the probability of a positive test result using the alcohol wash testing method varies with the total number of *varroa* in a hive. It is based on a single simulation instance of the model, looking at testing in a single hive with no movement of mites between neighbours. The probability of a positive test result using an alcohol wash is calculated using Eq. (5). This figure shows the probability of a positive test result oscillating as *varroa* transition from a phoretic state to being in the brood. The peaks in probability of a positive test result align with peak numbers of phoretic mites. This indicates that our model results are sensitive to test timing parameters – time until testing starts and time between testing. If testing differs by only a few days it could be the difference between testing during a peak in probability of a positive test result and a trough in probability of a positive test result.

**Figure 2:**
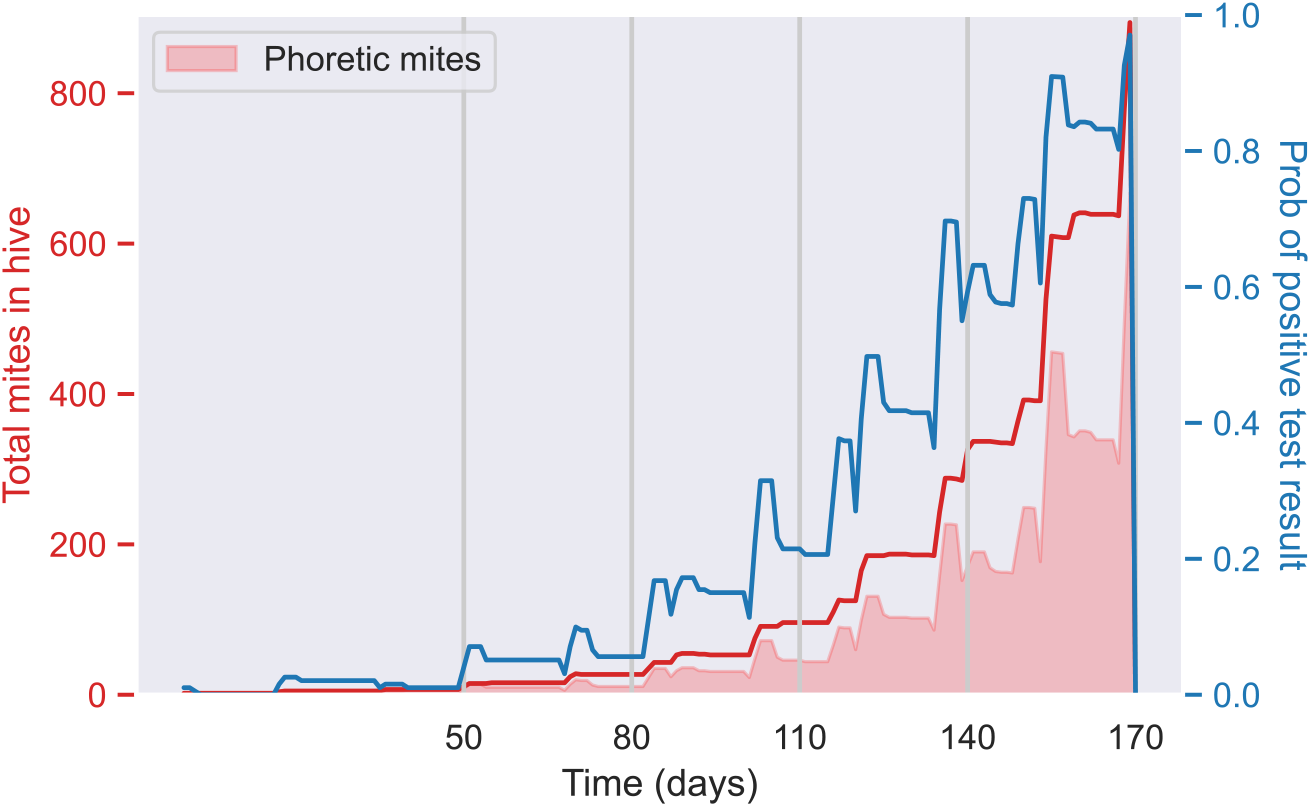
Total mites in a single hive and probability of a positive test result using the alcohol wash testing method over time (Eq. 5). The blue line indicates the probability of a positive test result through time, the red line indicates the total mites in the hive through time and the vertical grey lines indicate days when testing occurs. The red shaded area indicates the number of phoretic mites in the hive at a given time. We assume testing starts on day 50 and occurs every 30 days thereafter.

### 3.2 Emergent dynamics

In this study, we aim to mathematically model known biology of *varroa* reproduction to explore the spread of *varroa* in an outbreak setting. To provide context for later results, here we explore the dynamics of *varroa* spread between hives emerging from our modelling.

There are two competing stochastic elements at play in our model: stochastic death of *varroa*, either natural (Section 2.1.2) or through hive elimination (Section 2.1.4), and stochastic *varroa* movement between hives (Section 3.3.1). Figure 3 shows the two patterns of spread we see in our modelling: (a) *varroa* dies out/is caught before spreading to other hives, or (b) *varroa* spreads throughout the network before being caught/dying out in the initial hive. In the simulations where *varroa* are able to spread throughout the network, they most often spread extensively, infesting most, if not all hives on the network.

**Figure 3:**
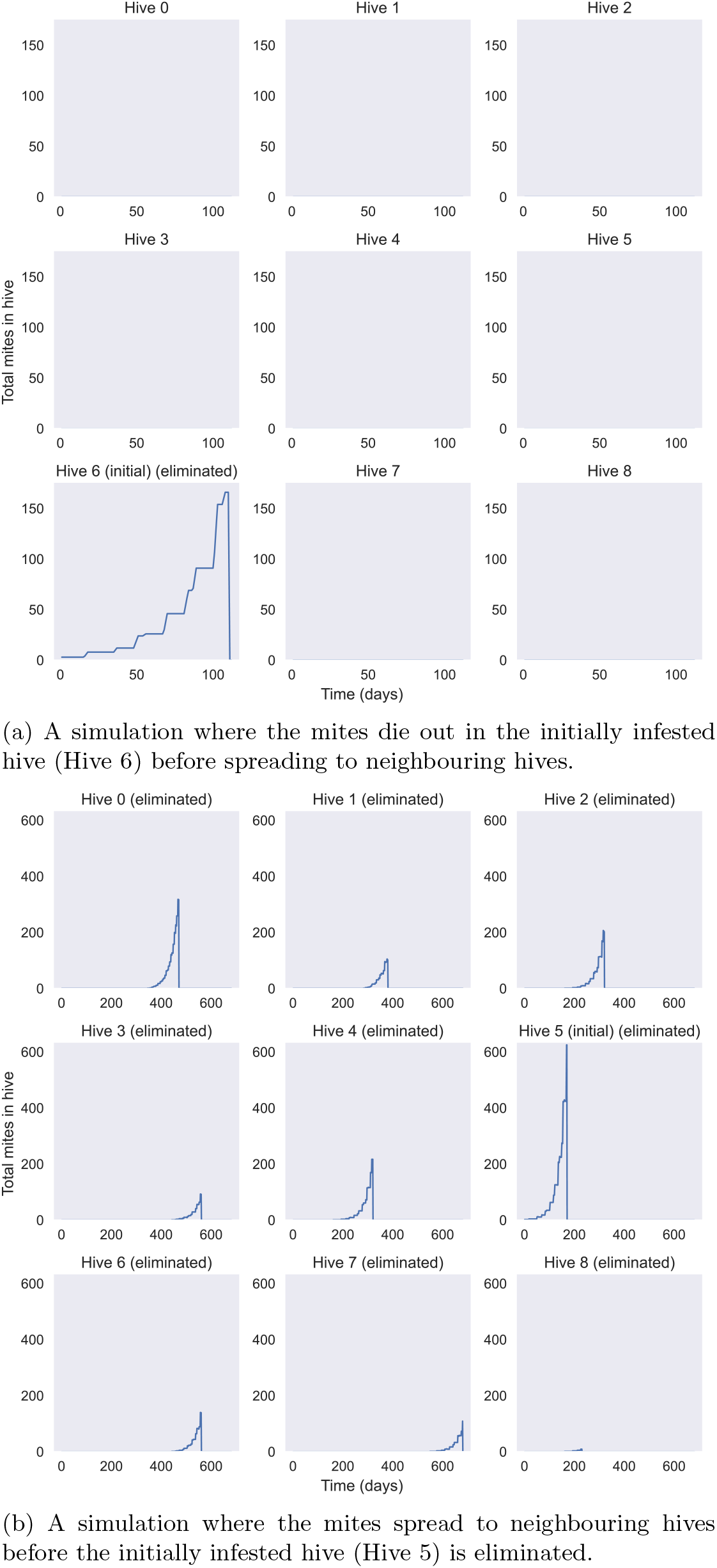
Single simulations showing two model outputs when we seed an infestation in a random hive: (a) *varroa* is eliminated before spreading further and (b) *varroa* spreads before the initial hive is eliminated. Each subplot shows the number of mites in each hive on the considered grid. The initially infested hive is denoted with “(initial)”. An eliminated hive is denoted with “(eliminated)”.

For the following analysis, we therefore see bimodal patterns. The first peak occurs around one hive eliminated – these are the simulations in which *varroa* has died out before spreading to neighbouring hives. The second peak occurs around a larger number of hives eliminated – these are the simulations in which *varroa* spreads through the network before being caught or dying out in the initial hive. Where this second peak is centred depends on the parameters considered.

### 3.3 Sensitivity analysis

Unless otherwise stated, the following results consider parameter values as described in Table 1. In Figures 4 – 8, we present the proportion of simulations where 1, 2 – 4, 5 – 8, or 9 hives are infested and subsequently eliminated as we vary different parameters. Note that we did not see any simulations where 0 hives were eliminated (i.e. the initial infestation died out before testing positive or spreading to neighbouring hives). As such the results presented range between 1 and 9 hives eliminated.

**Figure 4:**
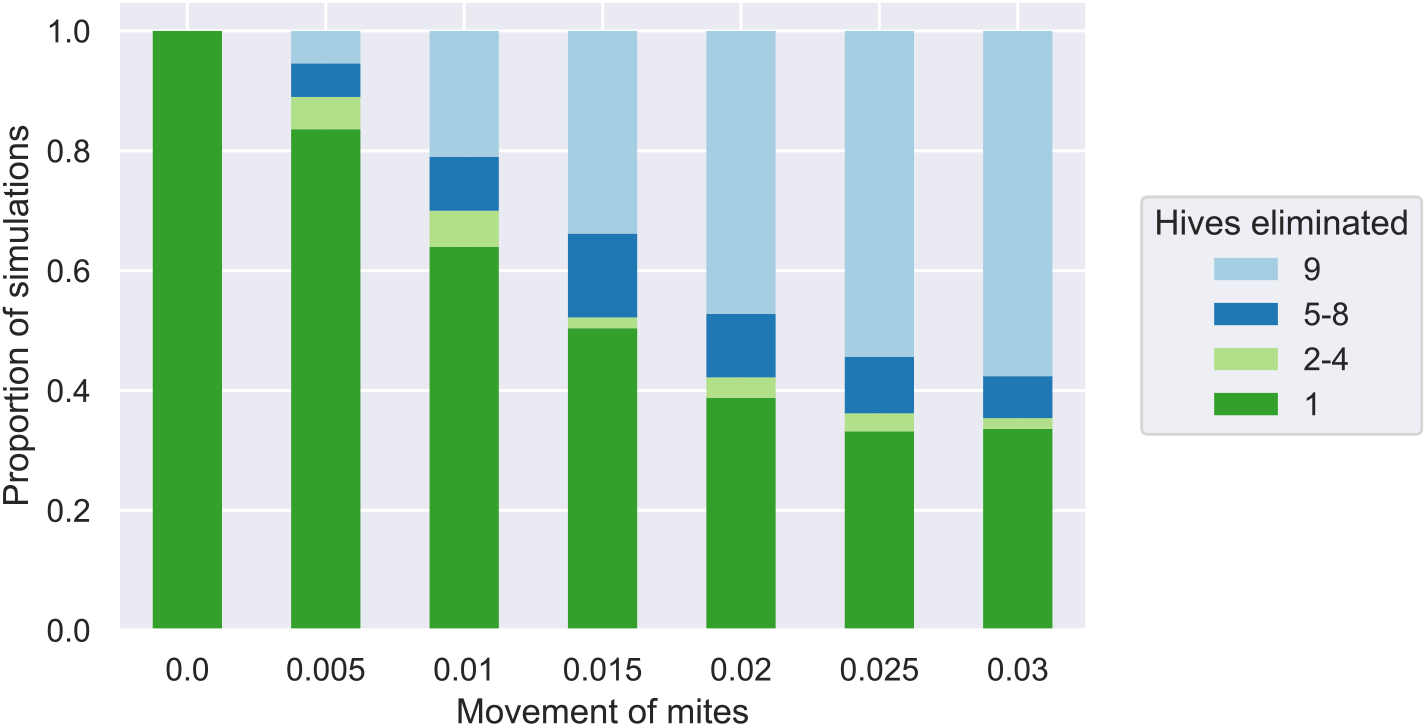
The proportion of simulations where 1, 2–4, 5–8, or 9 hives are eliminated as we vary the parameter representing the movement of *varroa* between hives (*α*). We consider 500 simulations on a grid of 9 hives for each parameter. The parameter considered, *α* represents the proportion of bees (and any mites on them) that leave a hive and subsequently move to a neighbouring hive (as described in Section 3.3.1).

In simulations with 1 hive eliminated, *varroa* did not spread from the initially infested hive. In simulations where 9 hives are eliminated, *varroa* spread to all hives in the network. Simulations with 2–4 hives eliminated indicate limited *varroa* spread through the network (minority of hives are infested and subsequently eliminated). Simulations with 5–8 hives eliminated indicate extensive *varroa* spread through the network (majority of hives infested and subsequently eliminated). Simulations where 1 or 9 hives are eliminated are displayed separately to the 2–4 and 5–8 groupings as together they make up the majority of simulations for all results.

#### 3.3.1 Movement between hives

Figure 4 demonstrates how the movement of mites between hives impacts the number of hives eliminated across 500 simulations. The movement parameter considered, *α*, represents the proportion of bees that leave a hive on a given day and migrate to a neighbouring hive (Eq. 8). For example, if we consider a movement parameter of *α* = 0.01, of the bees that leave a hive on a given day, 1% of those bees (and therefore also any *varroa* being carried by them) will end up in a neighbouring hive.

Increasing the movement of bees and any *varroa* they carry between hives increases the proportion of simulations where *varroa* spreads extensively before being caught and eliminated. When there is no movement of *varroa* between hives (*α* = 0), there can only be 1 hive eliminated (the initially infested hive). As we increase movement from *α* = 0.005 to *α* = 0.03, we see an increase in simulations with extensive spread of *varroa*. However, even in cases of increased movement, there still remain a proportion of simulations with limited/no spread where *varroa* is caught and eliminated before having a chance to spread. Increased movement, for instance due to hives being in close proximity for pollination services (e.g. (Feuvre 2018)), can potentially lead to extensive spread before *varroa* is caught by testing. In our modelling, restricting the movement of mites, in combination with testing and hive elimination, can reduce the proportion of simulations with extensive spread of *varroa* throughout a network of hives.

#### 3.3.2 Test timing

Figures 5, 6 and 7 demonstrate how the number of hives eliminated differs as we vary the time between testing, the test start day, and both the time between testing and test start day. As we saw in Figure 2, the number of hives eliminated will depend on the interplay between the time between testing and the test start day. As such, we expect the results to be sensitive to the parameters considered.

**Figure 5:**
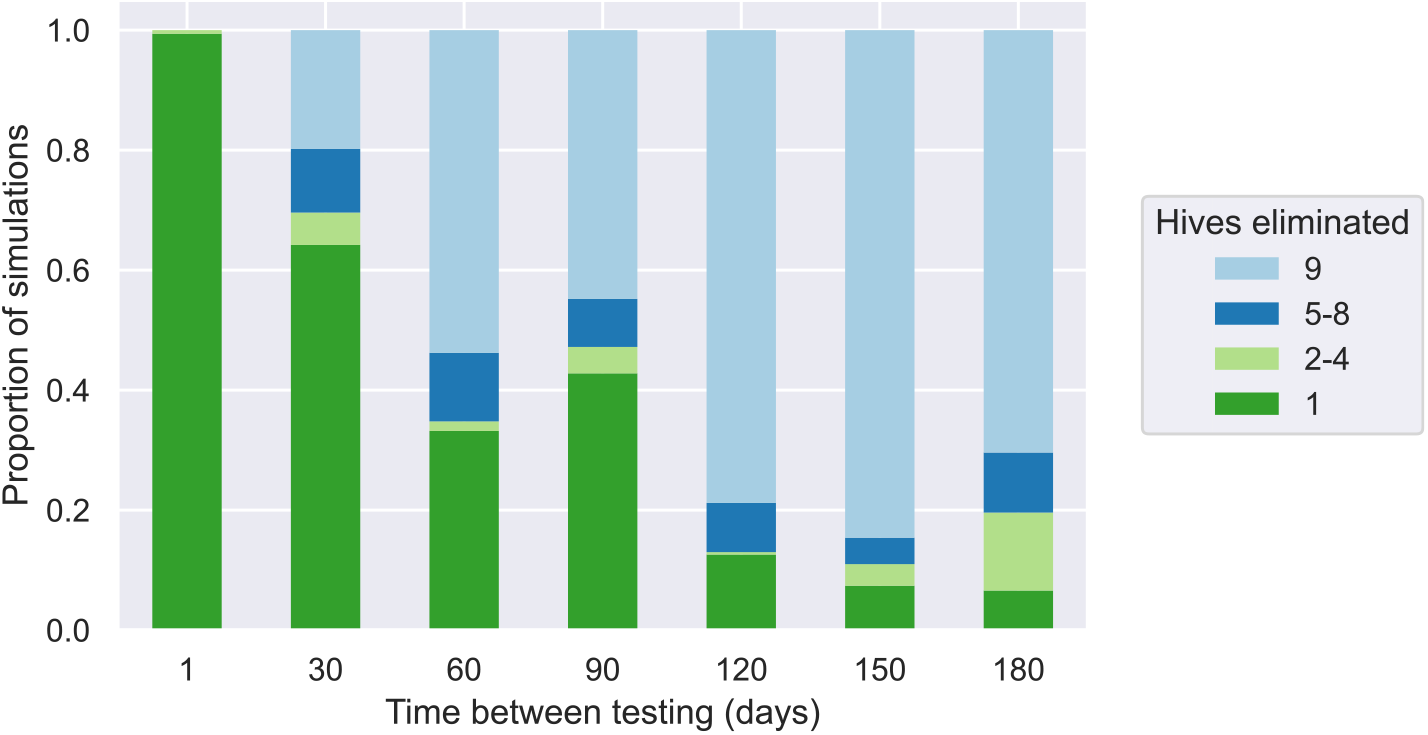
The proportion of simulations where 1, 2–4, 5–8, or 9 hives are eliminated as we vary the time between tests. We consider 500 simulations for a grid of 9 hives for each parameter. Testing starts at 50 days in each simulation.

**Figure 6:**
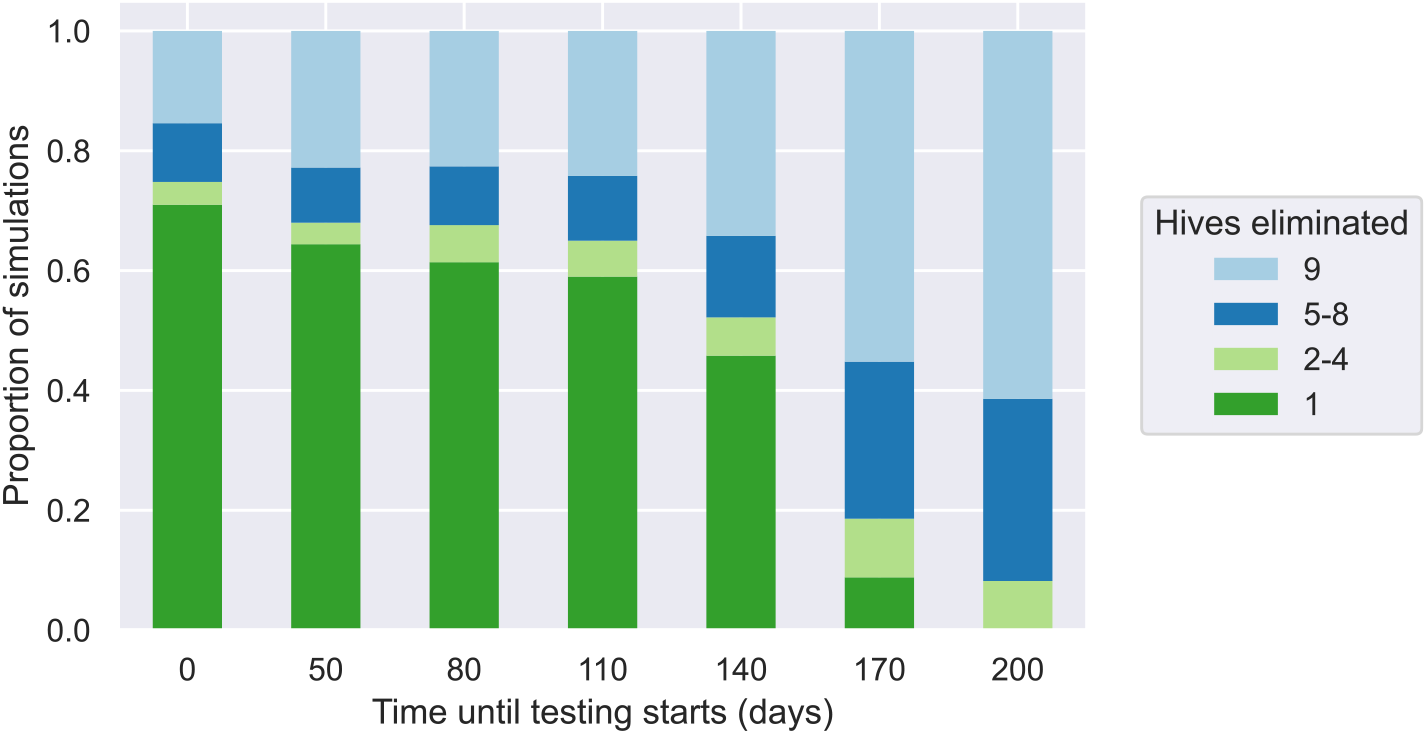
The proportion of simulations where 1, 2–4, 5–8, or 9 hives are eliminated as we vary the test start day. We consider 500 simulations for a grid of 9 hives for each parameter. The time between tests is fixed at 30 days to compare with results from Figure 5.

**Figure 7:**
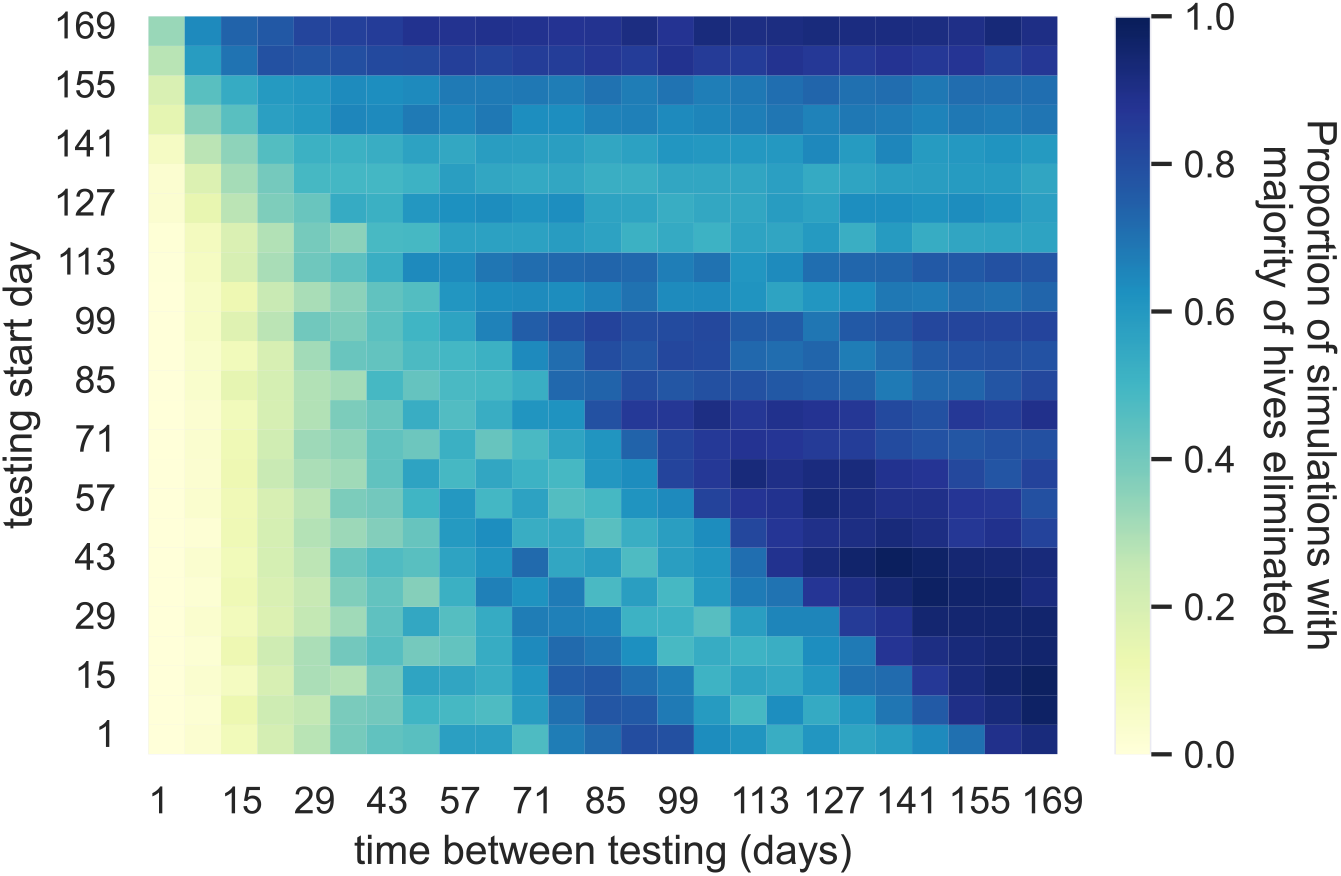
The proportion of simulations where a majority of hives are infested and subsequently eliminated (more than 4 hives eliminated) as we vary both the time between testing and the test start day. For each parameter combination we consider 500 simulations for a grid of 9 hives.

Figure 5 demonstrates how the number of hives eliminated varies as we change the time between testing across 500 simulations. As we increase the time between testing, we generally see an increase in simulations where *varroa* spreads extensively before being caught. At the extremes, when testing occurs every day (1 day between testing), *varroa* is caught in the initially infested hive before it can spread further in all simulations (1 hive eliminated for all simulations). Conversely, when testing occurs every 180 days we see that the majority of simulations have nine hives (all hives) eliminated. However, even when testing every 180 days there remain some simulations where *varroa* does not spread further than the initially infested hive, either through early detection and elimination or, by chance, there is no movement of *varroa* between hives before elimination.

However, Figure 5 also shows that testing more frequently does not necessarily correspond to fewer simulations with extensive spread of *varroa* for intermediate times between testing. Take for example the results when testing every 60 days compared to every 90 days. When testing more frequently (every 60 days) we see *more* simulations where *varroa* has spread extensively compared to testing less frequently (every 90 days). This indicates that although we are testing more frequently, hives are not testing positive for the first few rounds of testing. As seen in Figure 2, the probability of a positive test result, while oscillating depending on the number of phoretic mites in a hive, remains low while the *varroa* population is small. The peaks in probability of a positive test results also do not increase monotonically in time. Furthermore, there is a lower chance of movement of *varroa* between hives when the *varroa* population is small. Thus, testing more frequently when the *varroa* population is small does not give any advantage. It will take multiple rounds of testing before the *varroa* infestation has grown large enough to be detected by the tests. In general, testing more frequently does not guarantee reduced spread of *varroa*, particularly at the start of an outbreak when *varroa* infestations are small.

Figure 6 demonstrates how the number of hives eliminated changes as we vary the time until testing starts. We have chosen testing start day parameters to align with the testing schedule when considering 30 days between testing in Figure 5. That is, we demonstrate the effect of delaying testing from starting at day 50 to starting at day 80 or 110 etc. on the hives eliminated. The parameter value 0 has been added for comparison and represents testing from the first incursion and every 30 days thereafter. The second bar of Figure 5 and the second bar of Figure 6 show results using the same parameters (30 days between testing, testing starts at day 50), and any differences in results are due to stochastic variation in the model.

For testing start days of 0, 50, 80 and 110, we see results similar to simulations where the time between testing is 30 days in Figure 5 with minor variations due to model stochasticity. This indicates that when testing every 30 days, a positive test result is often not returned until around 110 days, i.e. the third testing round (when testing starts at 50 days). In many simulations, testing before this returns a negative test result. When testing starts after this (i.e. 140 days onwards), we see more simulations where the mites spread before testing. That is, if testing had started earlier, the mites could have been caught and further spread prevented. In particular, when we consider a start date of 170 or 200 days, the mites spread extensively before being caught in the vast majority of, or all, simulations. When varying the test start day, we alter the alignment of the testing schedule with the *varroa* reproduction cycle. The probability of a positive test result on a given day varies according to how the testing schedule aligns with the mite reproduction cycle in addition to the total mite population. Thus, starting testing earlier does not guarantee early detection.

Figure 7 demonstrates how the number of hives eliminated changes as we vary both the time until testing starts and time between testing. Testing frequently (1 – 15 days between testing) means *varroa* is detected before extensive spread for most testing start days considered. It is only as we approach start days of 141, 148 and 155 that we start to see an increase in the proportion of simulations with a majority of hives eliminated with frequent testing. Testing frequently helps overcome the limitations of our chosen testing methods: low probability of detection with small *varroa* populations. Even when testing is delayed, limited spread occurs before testing begins and *varroa* is detected shortly thereafter.

For more than 29 days between testing, as we hold the time between testing constant and increase the testing start day we see varying results for the proportion of simulations with majority of hives eliminated. These dynamics are most evident around 5 - 8 weeks between testing. As shown in Figure 2, the probability of positive detection varies as *varroa* go from a phoretic stage to reproducing in the brood, where they are undetectable by testing. As we increase the testing start day, the alignment of the testing schedule with the *varroa* reproduction cycle changes. This therefore alters how quickly *varroa* is caught by testing, impacting their potential to spread through the network of hives.

Testing early but infrequently produces similar results to testing late but frequently. As seen in Figures 5 and 6, early tests are often negative due to small *varroa* populations. Therefore, when testing early but infrequently, it is likely that only later tests will return positive detection. This allows *varroa* to spread extensively in between tests until infestations are eventually caught. When testing occurs frequently, but starts late, it is likely that *varroa* will be caught in an early test round. However, the mite population has had time to spread extensively before testing begins.

Testing late and infrequently does not necessarily result in guaranteed extensive mite spread. By the time testing starts, we expect there to be large enough *varroa* populations to be detected by testing. If *varroa* has not managed to spread throughout the network before the first test, an extensive incursion can be averted even when testing late. However, as we increase the time to the first test and the time between testing, we see the mites are able to spread extensively before being caught, even though this is likely to occur in the first testing round.

#### 3.3.3 Mites removed by testing

Figure 8a shows the number of hives eliminated across simulations as we vary the proportion of mites on sampled bees removed by testing. Results for the proportion of mites removed during sugar shake testing (*r* = 0.4) and alcohol wash testing (*r* = 0.7) are highlighted in Figure 8b. The proportion of mites on sampled bees removed by testing can be thought of as a type of test sensitivity. In our model, we assume any mites detected are *varroa* (i.e. 100% test specificity).

**Figure 8:**
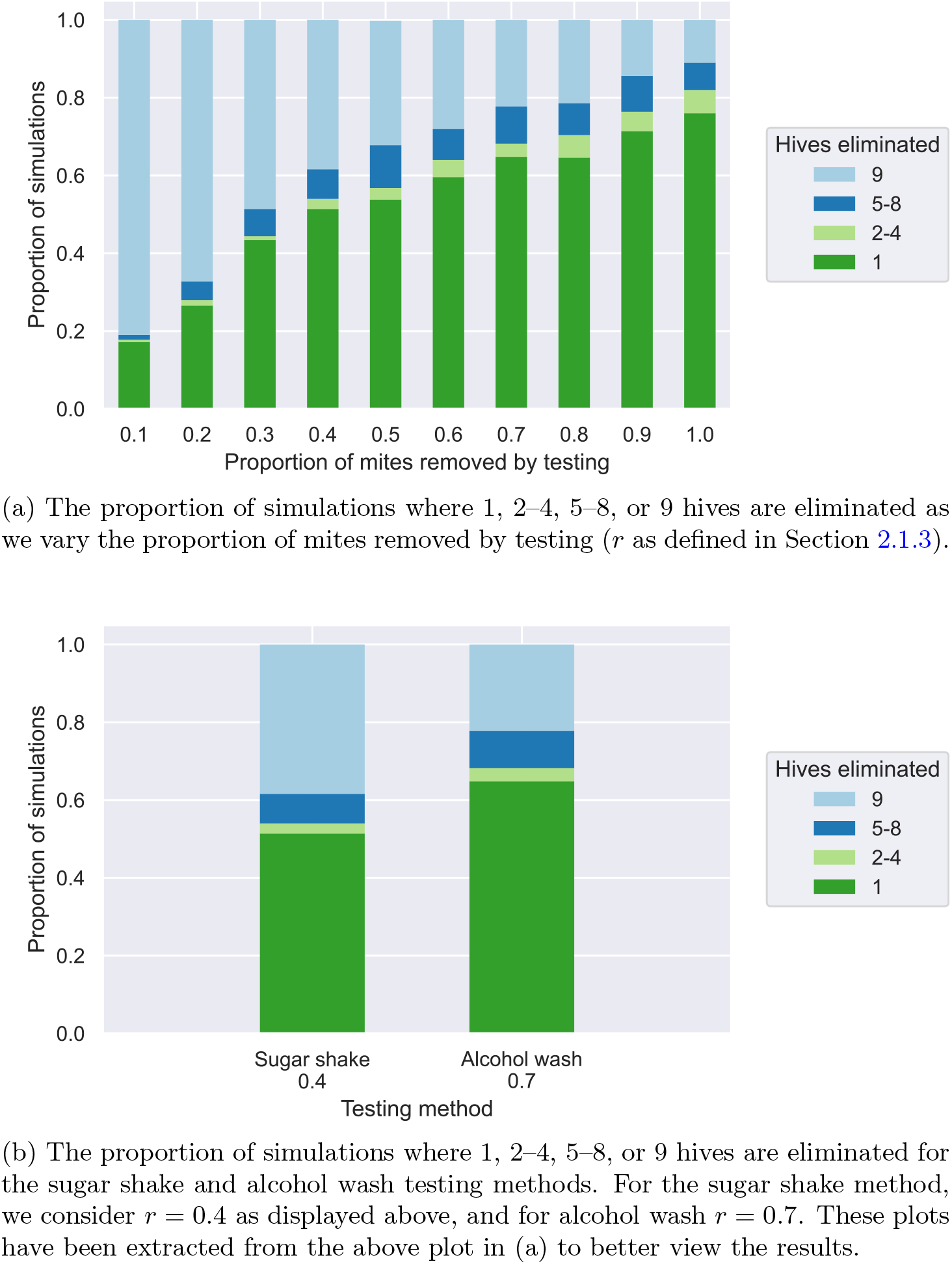
Sensitivity analysis on the proportion of mites removed by testing. For each parameter considered, 500 simulations for a grid of 9 hives are plotted.

For these results, we have considered testing starting 50 days after the initial incursion, 30 days between testing, and a movement parameter of *α* = 0.01. Given these parameters, we see when the proportion of mites removed by testing is increased, more simulations have lower numbers of hives eliminated. That is, there are more simulations where the outbreak is controlled before extensive spread of *varroa* through the network of hives. As the alcohol wash testing method removes a higher proportion of mites from sampled bees, we see a decrease in simulations with extensive spread compared to the sugar shake testing method. However, for all proportions considered, we still see some simulations where the mites have either died out or are caught before spreading further and some simulations where the mites spread extensively before being caught by testing and eliminated. In particular, we still see this for the extreme values of the removal proportion, i.e. very low proportion of mites on sampled bees removed by testing (*r* = 0.1) and all mites removed by testing (*r* = 1). When testing recovers almost no mites on sampled bees (*r* = 0.1) there is still potential for an incursion to be caught before spreading to neighbouring hives. When testing removes all mites on sampled bees (*r* = 1), there is no guarantee that the sampled bees carry any *varroa*. Thus, even when testing removes all mites on sampled bees, *varroa* can still potentially spread through the network before being detected.

These results are highly dependent on the test start date, time between testing and movement parameters considered. As shown previously, as we delay the test start date and/or increase the time between testing we expect mites to be able to spread further throughout the network before being caught by testing. Increasing the movement between hives will also result in increased chance of extensive spread before elimination. While the results in Figures 8a and 8b demonstrate how increasing the proportion of mites removed by testing (e.g. through using an alcohol wash instead of a sugar shake) can result in a higher chance of catching and eliminating an incursion before extensive spread, this can never be guaranteed.

## 4 Discussion

In this study, we present a mathematical model capturing the within-hive and between-hive dynamics of a *varroa* outbreak in a naive population of European honeybee hives. For our considered parameters, the majority of simulations end with either one or all hives infested and subsequently eliminated following an initial *varroa* incursion. Due to the considered testing strategies only capturing phoretic *varroa*, we see the probability of a positive test result vary as *varroa* move between being phoretic and being in brood. When varying movement between hives, we see that increased movement leads to an increase in simulations where a majority of hives are eliminated following infestation. Additionally, the dynamics of phoretic testing play an important role in the number of hives eliminated when varying test timing parameters, i.e. time between testing and time until testing starts. In these simulations, the number of hives eliminated greatly depends on how the testing schedule aligns with the lifecycle of *varroa* within hives. When testing does occur, there needs to be both a large enough *varroa* population and enough *varroa* in a phoretic state to ensure positive detection and elimination before *varroa* can spread to neighbouring hives. As the proportion of mites removed from sampled bees by testing increases, we see an increase in the proportion of simulations where a minority of hives are infested and subsequently eliminated following infestation. However, even when all mites are removed by testing (*r* = 1), we still see simulations where the majority of hives become infested before being eliminated.

In our simulations, the number of hives eliminated in a *varroa* outbreak is greatly influenced by the mite lifecycle. Irregular testing schedules could help overcome the limitations of testing only phoretic *varroa* in a hive. Due to the oscillatory nature of the probability of positive detection (Figure 2), regular testing schedules can detect *varroa* earlier when they align with peaks in probability of positive detection, and later when they align with troughs. Our results show the difference between detecting *varroa* early or late can come down to a few weeks. By contrast, an irregular testing schedule could capture both peaks and troughs in the probability of positive detection in the same schedule. Testing schedules provide a powerful tool to overcome limitations of a particular testing method, for example increasing testing frequency can help overcome low test sensitivity (Abell et al. 2023). In our model, an irregular testing schedule could look like testing once a week for 3 weeks, and repeating this once every 3 months. In reality, we cannot reliably know when day one of an outbreak is, and so we need robust testing strategies, independent of when the testing strategy begins. Incorporating irregularities into hive testing schedules could help detect *varroa* earlier than a regular testing strategy, reducing spread to neighbouring hives.

Our modelling incorporates known *varroa* biology and hive testing practises to understand the dynamics of early outbreak spread of *varroa* between hives. In Australia, it is likely that *varroa* was not detected until 10–24 months after the first incursion (Department of Agriculture, Fisheries and Forestry 2011; Owen, Stevenson, and Scheerlinck 2021). In general, we are unlikely to know when the initial incursion occurs and we rely on positive detection in sentinel hives to trigger routine testing for *varroa*. In our modelling, we assume a *varroa*-naive population engaging in consistent, frequent surveillance of sentinel hives. While this scenario is no longer relevant for New South Wales, neighbouring states can currently be considered *varroa*-naive populations engaging in frequent surveillance. As *varroa* continues to spread throughout Australia, there is the possibility for beekeepers to replace colonies that have been eliminated following positive detection. Modelling on this longer timescale would need to incorporate hive replacement, which would allow further spread of the mite and potentially extend the length outbreak. Furthermore, while our modelling captures movement of *varroa* between hives, it does not capture the various causes of this movement. There are a number of ways by which *varroa* carried on bees can move between hives – e.g. through bees being accepted into a neighbouring hive on the way home or from a colony robbing a weak neighbouring hive of resources (Messan et al. 2017). We parameterise movement in our model through considering the mites on the proportion of bees that leave one hive and migrate to a neighbouring hive. This allows us to consider scenarios with increased movement between hives (such as through mass pollination events), and scenarios with decreased movement (such as biosecurity movement restrictions). However, there are limitations in how movement between hives is considered in our model, and therefore how the movement parameters can be interpreted. There is currently no data on the exact proportion of bees migrating between hives on a given day. Furthermore, our model does not consider the impact of distance between hives on this migration. Modelling to support future *varroa* outbreak response would need to incorporate data from previous outbreaks to ensure model parameters and therefore results are suitable for a specific scenario.

When modelling real world testing practices, we need to be cognisant of the difference between a recommended practice and what realistically occurs. Generally, people are reluctant to opt for a testing method that kills bees over one that does not, i.e. alcohol wash testing versus sugar shake testing, without adequate justification for the former. Testing strategies need to consider individual impacts on beekeepers, particularly with strategies that recommend eliminating hives on positive detection. Strategies lacking justification could create a disincentive to test and subsequently report infestations, potentially leading to further spread of an outbreak (Hernández-Jover et al. 2016; Palmer, Sully, and Fozdar 2009). Unlike other countries, European honeybees only overwinter in the coldest areas of Australia, a behaviour that kills off *varroa* in a hive over winter months (Goodman 2014). As such, Australian beekeepers need to monitor and manage *varroa* incursions year-round. With the threat of national *varroa* spreading nationally imminent, Australia needs to develop comprehensive testing and management strategies to protect beekeepers and their hives. Designing successful testing strategies requires connecting mathematically robust strategies with realistically feasible practices that can be replicated in real life. This work presents a first step in utilising mathematical modelling to understand the impact of testing and elimination strategies on outbreak dynamics.

## Supplementary material

The supplementary material, code and data for this paper can be found at https://github.com/iabell/bee-all-and-end-all.

## Acknowledgements

The authors would like to acknowledge Greg Chandler for their invaluable advice and support for this work. This work was conducted on the lands of the Wurundjeri people of the Kulin nation.

